# Non-negative Independent Factor Analysis disentangles discrete and continuous sources of variation in scRNA-seq data

**DOI:** 10.1101/2020.01.31.927921

**Authors:** Weiguang Mao, Maziyar Baran Pouyan, Dennis Kostka, Maria Chikina

## Abstract

**Motivation:** Single-cell RNA-seq analysis has emerged as a powerful tool for understanding inter-cellular heterogeneity. Due to the inherent noise of the data, computational techniques often rely on dimensionality reduction (DR) as both a pre-processing step and an analysis tool. Ideally, dimensionality reduction should preserve the biological information while discarding the noise. However if the dimensionality reduction is to be used directly to gain biological insight it must also be *interpretable* – that is the individual dimensions of the reduction should correspond to specific biological variables such as cell-type identity or pathway activity. Maximizing biological interpretability necessitates making assumption about the data structures and the choice of the model is critical.

**Results:** We present a new probabilistic single-cell factor analysis model, **N**on-negative **I**ndependent **F**actor **A**nalysis (NIFA), that incorporates different interpretability inducing assumptions into a single modeling framework. The key advantage of our NIFA model is that it simultaneously models uni- and multi-modal latent factors, and thus isolates discrete cell-type identity and continuous pathway activity into separate components. We apply our approach to a range of datasets where cell-type identity is known, and we show that NIFA-derived factors outperform results from ICA, PCA, NMF and scCoGAPS (an NMF method designed for single-cell data) in terms of disentangling biological sources of variation. Studying an immunotherapy dataset in detail, we show that NIFA is able to reproduce and refine previous findings in a single analysis framework and enables the discovery of new clinically relevant cell states.

**Availability:** NFIA is a R package which is freely available at GitHub (https://github.com/wgmao/NIFA).

**Supplementary information:** Supplementary data are available at *Bioinformatics* online.

## 1 Introduction

Single-cell RNA sequencing (scRNA-seq) techniques have allowed researchers to query the complexity of transcription regulation at an unprecedented level of detail. scRNA-seq technologies have the power to reveal both distinct cell types and transcriptional heterogeneity within a defined cell population. As the the number and size of single-cell datasets increases, it becomes important to develop methods that can quickly summarize the *biological* information embedded in a scRNA-seq dataset as a set of interpretable variables that can be used for downstream analysis. One kind of summary measure is the identity and number of cell types present in a dataset. In recent years there has been a proliferation of clustering methods designed to address this problem (Kiselev *et al.,* 2017; Butler et *al.,* 2018). Clustering approaches assume that the data is well described by a discrete set of cell types, but in many cases, questions about continuous biological variation, such as developmental trajectories or levels of pathway activation are also of interest.

Such continuous variables do not conform to the assumptions of clustering algorithms but can be effectively modeled as latent factors. For example, cell-cycle variation has been repeatedly discovered in single-cell data, both using sophisticated latent variable models (Buettner *et al.,* 2015) and simple Principle Component Analysis (PCA)(Kowalczyk *et al.,* 2015).

Of course, cell-type/cluster identity can also be thought of as a latent factor, albeit a discrete one, thus suggesting that it is possible to design a single factor analysis framework that can discover both cell types and continuous biological variables in a single analytic step. In the ideal case a factor analysis representation of a scRNA-seq dataset would identify both discrete cell types and continuous pathway effects and importantly maximally disentangle difference sources of biological variation. In other words each factor should represent a single biological variable that could be measured directly using a different assay.

To address these goals we propose **N**on-negative **I**ndependent **F**actor **A**nalysis (NIFA) that can simultaneously models uni- and multi-modal factors thus combining clustering and latent variable analysis into a single framework. Our modeling framework is optimized to induce disentanglement between biological sources of variation (such as different cell types and pathways) so that individual factors are maximally aligned with specific biological variables and thus can be used in the downstream analysis. We evaluate our method on simulated and real data (see Table S1), and we show how it can be used to glean new biological insights by detailed analysis of an immunotherapy scRNA-seq dataset (Sade-Feldman *et al.,* 2018).

## 2 Methods

### The statistical model

*X* represents the RNA-seq matrix with dimension *P*-by-*N*, *A* denotes loading matrix with dimension *P*-by-*K*, *S* stands for latent variables with dimension *K*-by-*N*. We denote the *n_th_* column of *X* as *X_n_* = (*X*_1*n*_, *X*_2*n*_,…,*X_Pn_*)^*T*^, the *j_th_* row of *A* as 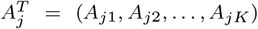 and the *n_th_* column of *S* as *S_n_* = (*S*_1*n*_, *S*_2*n*_,…, *S_Kn_*)^*T*^ (see Figure 1A and Supplementary Methods for details).

**Fig. 1.**
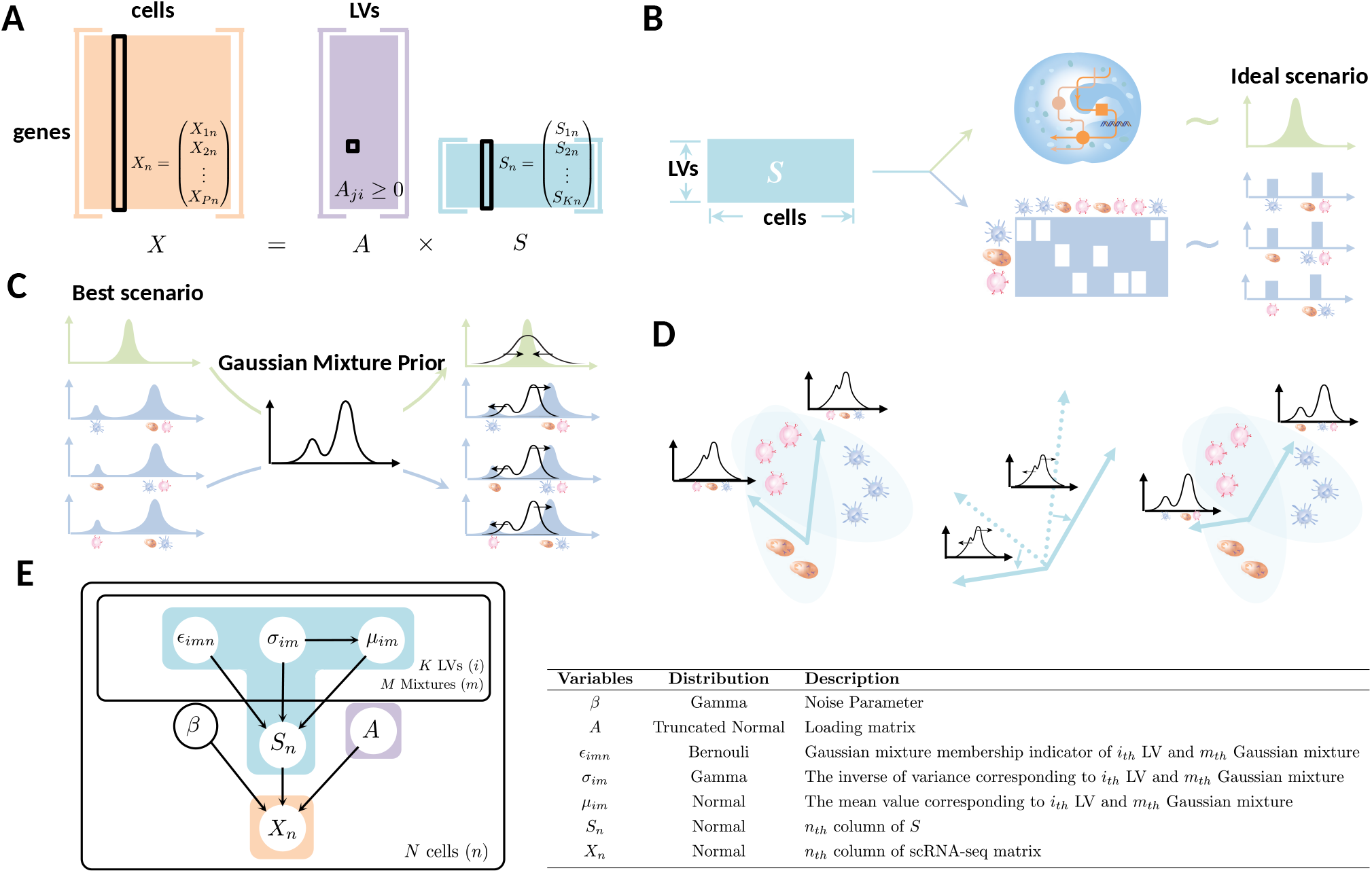
(A)NIFA decomposes the scRNA-seq matrix *X* into loading matrix *A* and latent variable matrix *S*. The loading matrix *A* is constrained to be non-negative. (B)NIFA can disentangle the discrete and continuous variations into different latent variables. In the ideal scenario, the underlying distribution of latent variables related to pathway effects should be unimodal while the underlying distribution of celltype-related latent variables follow bimodal distributions with clear separation between modes. (C) Both unimodal and multimodal variations can be jointly modeled by Gaussian mixture prior. The number of modes can be determined automatically via inference. (D) By imposing multi-modal prior, we force the latent factors to rotate and align with the directions that can best separate the cell-type identity. (E) Overview of the NIFA probabilistic graphical model. The table summarizes the probabilistic variables and the assumption of prior distributions. The details can be found in the Supplementary Methods.

For the factor matrix *S* we assume for each row that entries are iid samples from a (row/factor-specific) Gaussian mixture distribution with *M* components, parameterized by mean *μ_im_* and variance *σ_im_* (for row *i* and component *m*). Let *ϵ_imn_* (*i* = 1,…,*K*, *m* = 1,…, *M*, *n* = 1,…, *N*) be an indicator variable with *ϵ_ijn_* = 1 if *S_in_* was generated by component *j* and zero otherwise. Then we get for the prior of the *n*-th column of *S*:

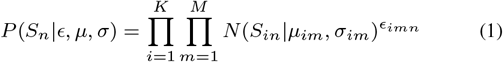

The loading matrix *A* is modelled with a truncated normal prior where *a* = 0 and *b* = ∞ indicating each entry *A_ji_* falls within the interval [0, ∞). *η* and λ denotes the mean and the inverse of the variance. Φ(·) is the cumulative distribution function of the standard normal distribution.

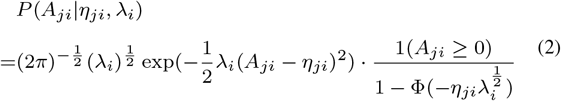

We assume the noise model to be Gaussian with a single precision parameter 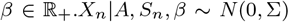 with Σ^-1^ = diag(*β*). Thus 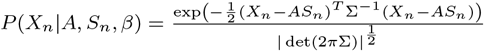.

### Joint likelihood & Parameter inference

The joint likelihood *P*(*X, A, S, e, μ, σ, β*) is as follows (Eq. 3).

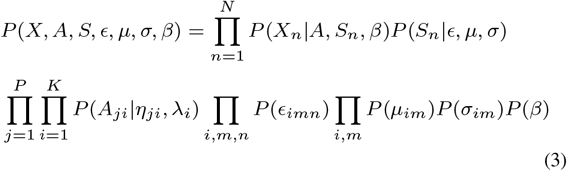

In order to efficiently infer the parameters, we apply variational inference technique, more specifically, mean-field approximation (Blei *et al.,* 2017). By assuming each variational parameter is independent of each other, we formulate the joint posterior distribution *Q*(*S, A, e, μ,σ, β*) (see Eq. 4) for the model and minimize the KL-divergence between Eq.3 and Eq.4 to derive the expression *q*(·) for each variational parameter as an approximation of single posterior distribution.

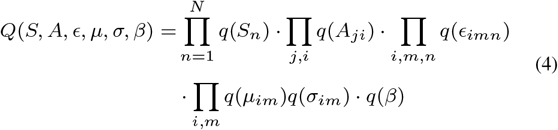

### Hyperparameter tuning

Our model has a number of hyperparameters, however, most of them are Bayesian priors and have relatively little impact on the results. One of the main hyperparameters of considerable relevance is the number of latent factors *K*. For the factor analysis model, the typical approach is to calculate the likelihood or ELBO (variation lower bound), comparing values directly or with BIC criterion (Krumsiek *et al.,* 2012) and selecting *K* corresponding to the optimal values. Since scRNA-seq data often has thousands of cells, the computation for likelihood-based or ELBO-based tuning method is time-intensive and impractical. Instead we can rely on variance-based method SVD with BIC criterion (Allen *et al.,* 2014) to figure out a conservative estimate of the number of latent factors. Importantly we use this only as a reference value, we perform all our evaluations across a range of *K* parameters.

We have one more discrete hyperparameter which is the number of Gaussian mixtures *M* for each latent factor. However, this parameter just needs to be set as the maximum number of components one can expect to find. Since our model fits the Gaussian mixtures by variational inference, it has the desirable property that *M* can be determined automatically as some of the mixing coefficients go to 0. In experiment we set this hyperparameter to be 4 and find that for non-developmental datasets, where we expect to find discrete cell types, the final number of modes is usually either one or two. This conforms to the biological intuition that cell types differ from each other by a set of (not necessarily unique) “marker” genes. Such marker genes typically have a bimodal distribution corresponding to high and low expression. While the distributions may overlap due to technical noise, we typically do not observe intermediate expression modes.

### Availability of data and code

We evaluate NIFA on several “gold” or “silver standard” scRNA-seq datasets (Gong *et al.,* 2018) with pre-annotated cell types. Table S1 summarizes the datasets we use. Gold standard datasets contain relatively homogeneous cell lines and experimental conditions are well controlled, while silver standard datasets define cell types based on expert knowledge. Most data were obtained through (https://github.com/hemberg-lab/scRNA.seq.datasets), or through corresponding GEO repositories: GSE120575 (Sade-Feldman *et al.,* 2018), GSE133656 (Liu *et al.,* 2019), and GSE70245 (Olsson *et al.,* 2016). We also include simulated datasets generated by Splatter (Zappia *et al.,* 2017), using Kumar (Kumar *et al.,* 2014) and Zheng (Zheng *et al.,* 2017) as simulation input (Duò *et al.,* 2018).

## 3 Results

### Non-negative Independent Factor Analysis (NIFA): Theory

NIFA is a probabilistic matrix factorization model which decomposes the scRNA-seq matrix *X* into the loading matrix *A* and the sources/latent variables *S* (Figure 1A). While factor analysis models are routinely used for pre-processing, if the factors themselves are to be analysed it is important that they are interpretable, that is they should have a clear correspondence to a specific biological variable. A general popular approach to enhance interpretability is enforcing certain structures or constrains for the latent variables or their loadings such as positivity (as exemplified by the successful non-negative matrix factorization (NMF) and variants (Stein-O’Brien *et al.,* 2019; Kotliar *et al.,* 2019; Zhu *et al.,* 2017; Sherman *et al*., 2020)). NIFA maximizes interpretability by explicitly modeling the generative process of single cell data as a combination of two types of variables: cell-type identity and pathway activity.

While the semantic difference between these is not absolute, we can differentiate them in terms of their statistical properties. For the purpose of our model we assume that cell identity corresponds to clusters, which in a generative model would correspond to discrete variables. Conversely, pathway activity correspond to continuous variation, with cell cycle being a canonical example. NIFA is designed to optimize the interpretability by isolating these factors into separate components. Figure 1B corresponds to the ideal representation of the discrete (depicted in blue) and continuous (depicted in green) generative variables. Typically discrete latent variables are inferred via clustering while continuous variables are inferred via matrix factorization/ factor analysis.

The key contribution of NIFA is that it unifies both procedures into a single framework. To do this the NIFA model considers the continuous relaxation of clustering, where clusters correspond to multimodal continuous instead of discrete variables (Figure 1C, blue factors). Each factor is modeled with a mixture of Gaussians prior which encourages multi-modal factors to become concentrated around their modes, thus enhancing the separation between high density regions (Figure 1C). The multi-modality is maximized when the factor is optimally aligned with a single cluster and thus the model encourages factors that represent *individual cell types* (Figure 1D).

This procedure is similar to the popular technique of Independent Component Analysis (ICA) (Hyvärinen and Oja, 2000), a DR technique which instead of attempting to model all the variation in the data finds maximally non-Gaussian components. ICA is a popular single-cell DR technique because it finds directions that maximally separate different cell types (Butler *et al.,* 2018). One important difference between ICA and NIFA is that for NIFA the exact form of non-Gaussianality is explicitly specified as Gaussian mixtures, whereas ICA more general.

A second important difference between ICA and NIFA is that NIFA doesn’t prefer multi-modal factors but only “enhances” them if it finds them. This flexibility arises from the Bayesian updates which allow the model to automatically determine the number of Gaussian mixture components for each factor. Since in the model factors are encouraged to concentrated at their modes, multi-modal components become “more multi-modal” while the shape of the uni-modal components is changed relatively little (see Methods for more details). It is important to note that since the Gaussian mixture priors reduce to a single Gaussian in the unimodal case the mixture assumption cannot contribute significantly to the interpretability of uni-modal factors. However, like NMF the NIFA model includes a non-negativity constraint on the factor loadings (Figure 1E) which encourages parsimonious and biologically interpretable factors. Thus, for the case of uni-modal factors NIFA defaults to NMF-like behaviour. Overall, our NIFA model combines assumption about the factors and the loadings to effectively induce disentanglement between individual cell-type identity factors as well as continuous pathway activity.

### Simulation Studies

We simulate artificial data considering 2,000 genes (*P* = 2000), 500 cells (*N* = 500), six latent factors (*K* = 6) and two mixture components per factor (*M* = 2, see Figure 1A and Supplementary Methods for full details). Briefly, we simulate latent factors and their mixture components independently, but the columns of the loading matrix *A* are correlated (details in Supplementary Methods). For the decomposition, we set *K* = 8 (all methods) and *M* = 3(NIFA) to see if NIFA can robustly recover the right latent factors given larger *K* and *M*, which is often the case in practice. Figure 2 summarizes our findings. While no method recovers all latent factors correctly, NIFA is able to more accurately recover latent factors compared with other methods. Also, NIFA correctly recovers the number of mixtures per factor as two (reporting zero for one of three possible mixture components). We also explored the same setting as above but with uni-modal factors (*M* = 1) and find that in this case NIFA performs as well as ICA (parallel), and that both NIFA and ICA (parallel) outperform other methods (data not shown).

**Fig. 2.**
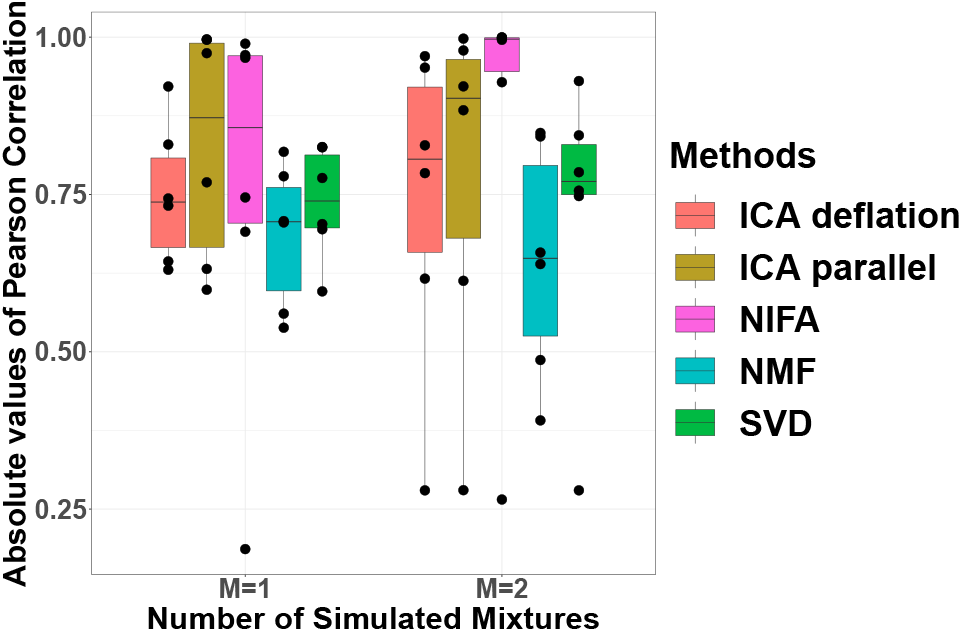
Evaluation on a simulated dataset. Boxplot of the correlation between simulated *S* and those recovered by SVD, ICA (parallel), ICA (deflation), NMF and NIFA. We find that the best performance is achieved by NIFA compared with all the other common methods.

### Evaluation

Meaningful evaluation of factor models is not necessarily straightforward, and several different options exist. A natural metric is the reconstruction error of a factor model (Levitin et al., 2019). However, for any specific decomposition there exist infinitely many alternative decompositions with exactly the same reconstruction error; yet, these may differ greatly with respect to the individual factors and loadings and, perhaps, biological implications.

Biological interpretability of decompositions is one of the attraction of factor models, but it is harder to evaluate and quantify than reconstruction error. Given substantial knowledge about the data is at hand, we can describe as a desirable property of an *interpretable* factor analysis is that there is *one-to-one* correspondence between inferred factors and known variables that significantly contribute to data variance. For example, factors that have one-to-one correspondence to known cell types are straightforward to interpret and can directly be used in subsequent analyses. However, if factors correspond to combinations of cell types (which could be an equally valid solution in terms of reconstruction error), interpretation is difficult and usefulness is more limited.

Therefore our evaluations are focused on assessing how well factor analysis models are able to disentangle known sources of biological variation. Such evaluations must necessarily reference some external ground truth, and for scRNA-seq data we consider two sources: known cell-type identity (used to evaluate factors) and gene to pathway membership (used to evaluate loadings). With this approach we compare the performance of NIFA with SVD, ICA, NMF and scCoGAPS (Stein-O’Brien *et al.,* 2019) across 14 data sets (see Table S1). scCoGAPS is a Bayesian NMF method that uses an ensemble-based approach to subset cells and collects consensus factors from parallelized Bayesian NMF runs across subsets.

#### Cell type identification

Here we investigate the ability of NIFA and five other methods to recover latent factors corresponding to pre-annotated cell types. In the case of scRNA-seq data the gold or silver standard of cell-type identity (see Table S1) is one such variable. In the ideal case each cell type corresponds to a unique factor in the model. In order to evaluate this property we compute maximum one-to-one correlations between factors and cell-type assignments (Figure 3). We find that on average our NIFA model performs better than ICA, NMF and scCoGAPS at this cell type detection task.

**Fig. 3.**
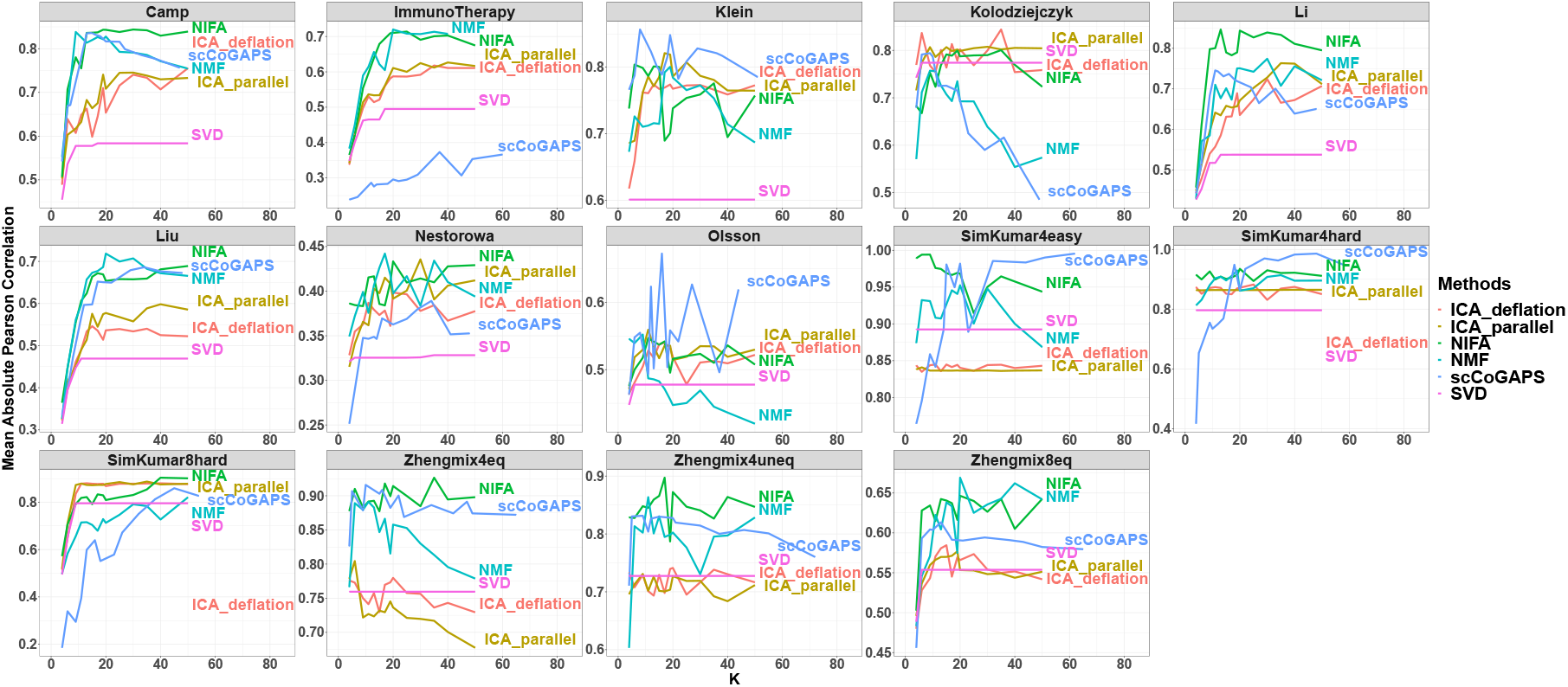
Evaluation of one-to-one correspondence between factors and cell types. Given a set of factors and a set of cell-type labels, we evaluate the maximum correlation between each cell type and a factor. For clarity, we plot the mean correlation values across all cell types. In order to account for the possibility that different models may need different number of factors (*K*), we report the results at varying *K*. We compare NIFA with ICA, NMF (KL-loss), scCoGAPS and SVD as a baseline. We find that on average our NIFA model performs better than competing decomposition methods on average and importantly while there can be large differences among NMF, ICA, the performance of NIFA (which combines features of both) always keeps track with the best method.

Importantly, while there can be large differences between NMF and ICA, the performance of NIFA (which combines features of both) always tracks with the best method.

#### Pathway enrichment

One important feature of factor analysis models is that the factors should be interpretable even in the absence of any ground truth knowledge. In such cases the factors are interpreted by inspecting the genes in their loadings. The expectation is that for a factor that captures a unique biological variable (which could be binary cell type or continuous pathway activation), the top loading genes are enriched for a few known functional modules. We evaluate this property by computing pathway enrichment for each factor using a hypergeometric test with the top 500 genes as foreground. This evaluation strategy allows us to evaluate existence of a biological focus in the model, independent of cell-type annotation. In this way the model can be credited for finding factors which capture pathway or cell-type signals even if these do not correspond to an annotated cell type. The pathway databases we use are “canonical pathways” from MSigDB (Liberzon *et al.,* 2011) and a comprehensive set of cell-type markers from xCell (Aran *et al.,* 2017). For canonical pathways we excluded pathways that had greater than 20% overlap with ribosomal or mitochondria genesets (defined as “KEGG_RIBOSOME” and “KEGG_ OXIDATIVE_PHOSPHORYLATION” respectively). We found that these are consistently enriched but provide little biological insight as variation in these pathways is often technical.

We then quantify the overall biological enrichment of a single loading vector as the mean fold enrichment for pathways that are significant at FDR<0.05. Pathway enrichment metrics summarized across all factors are plotted in in Figure 4 and Figure 5 for canonical pathways and xCell respectively. We find that not surprisingly the performance of all methods is much better for real biological datasets than simulated ones (Zhengmix4eq, Zhengmix4uneq and Zhenmix8eq). We also find that among the biological datasets NIFA is a consistently top performer in both “canonical pathway” and xCell evaluations, though the effect is more dramatic for xCell.

**Fig. 4.**
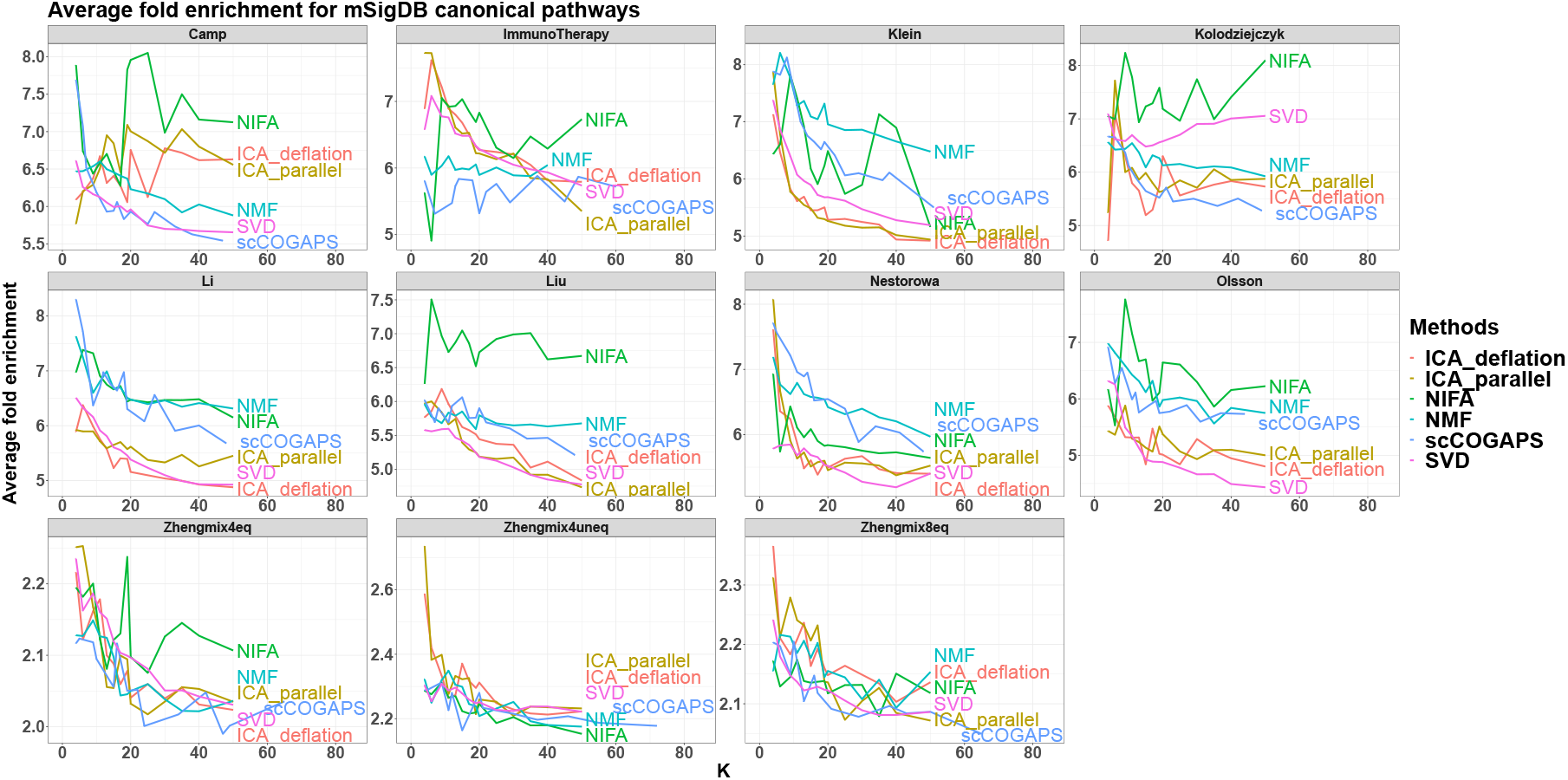
Pathway enrichment of “canonical pathways” from mSigDB (Liberzon et al., 2011). Enrichment is quantified as average fold enrichment among all factor-pathway pairs where the pathway is significantly over-represented in the top 500 loading genes (hypergeometric test, FDR<0.05). The first two rows are biological datasets. The last row (Zhengmix4eq, Zhengmix4uneq and Zhenmix8eq) corresponds to simulated datasets. All SimKumar datasets are excluded from this evaluation as they were not supplied with real gene names.

**Fig. 5.**
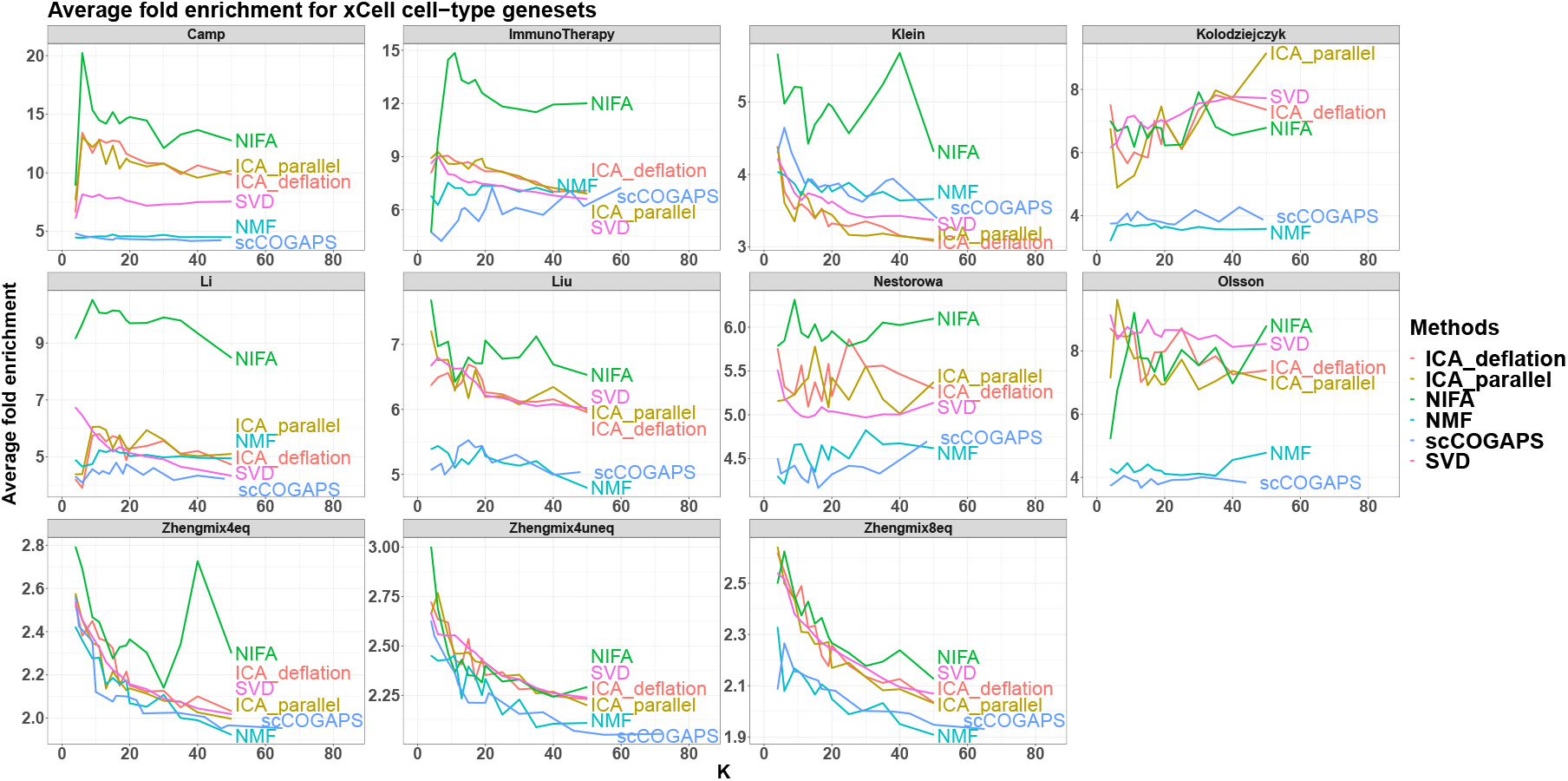
Pathway enrichment for xCell cell-type signatures. This figure is generated identically to Figure 4 but using the xCell genesets.

### In-depth evaluation of the Sade-Feldman *et al*. immunotherapy dataset

Antibodies that block immune checkpoint proteins, including CTLA4, PD-1, and PD-L1 are increasingly used to treat a variety of cancers. While checkpoint inhibitor (CI) therapy can be remarkable effective, not all patients respond (Larkin *et al.,* 2015). Determining the biological factors that facilitate or impede response to CIs remains an important and unresolved problem. In order to demonstrate how NIFA can be used to gain biological insight, we performed several in-depth analyses of the Sade-Feldman *et al*. immunotherapy dataset (Sade-Feldman *et al*., 2018). This dataset consists of 16,291 individual immune cells from 48 tumor samples of melanoma patients treated with checkpoint inhibitors. The dataset contains both pre-treatment and post-treatment samples and the patients are classified into responders and non-responders.

We applied our NIFA model to the entire single-cell dataset using *K*=25 which corresponds to the *k* with maximal correlation with known cell-type annotations (see Figure 3). The distributions of the inferred factors and the corresponding inferred Gaussian mixture fits are plotted in Figure S1. Each NIFA factor that has the best correspondence to existing annotations is given the same name. NIFA finds both uni- and multimodal factors and as expected the multi-modal factors are more likely to correspond to cell types.

In order to investigate which variables are associated with immunotherapy response, the resulting factors were mean aggregated to a single value for each unique patient sample. We also summarized the human-annotated celltype indicators as their mean values, corresponding to fraction of cells in sample. The resulting summary statistics were tested for association with response using Wilcoxon ranksum test and Benjamini-Hochberg FDR adjustment (separately for NIFA factors or human annotations). Pre- and post-treatment samples were analyzed separately and the resulting variables that were significant in either the pre-treatment or post-treatment comparison at FDR< 0.2 are plotted in Figure 6A. Top loading genes corresponding to each significant NIFA factor are show in Table 1.

**Fig. 6.**
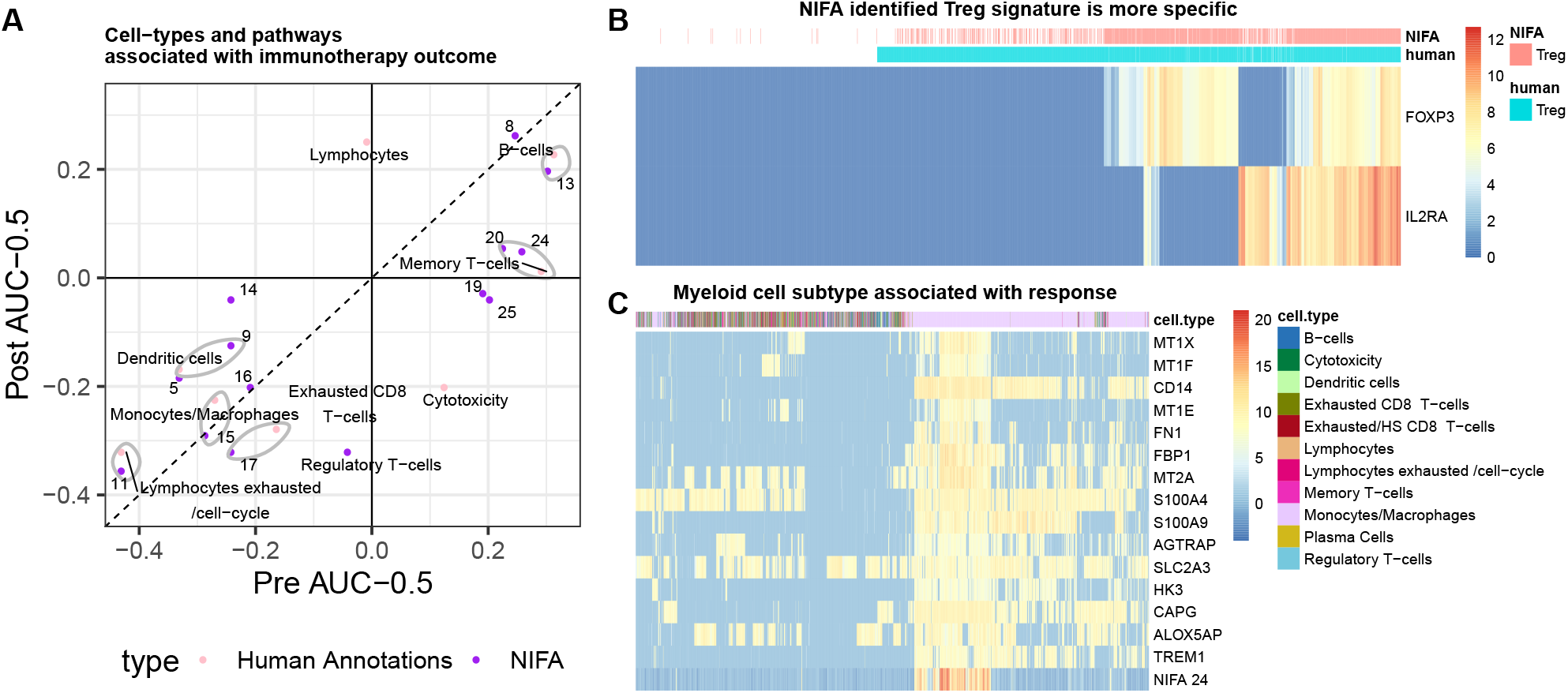
NIFA analysis of signatures associated with immunotherapy response. (A) NIFA-derived signatures of human-annotated cell types are mean aggregated per patient sample and the resulting summary statistics are tested for association with immunotherapy response. Variables that are significant at FDR<0.2 are shown with their respective normalized and centered ranksum statistics (ranksum/(number-of-positives × number-of-negatives)-0.5, equivalent to binary classification AUC-0.5). Pre-treatment effects are on the x-axis and post-treatment effects are on the y-axis. NIFA variables most closely matched to human annotations are grouped with grey ellipses. (B) Differences in Treg (Regulatory T-cells) identification between NIFA and human annotations. Heatmap of canonical Treg marker genes (FOXP3 and IL2RA) across all cells annotated as Tregs by either method and 1,000 randomly sampled other T cells. Overall, NIFA identifies fewer Treg cells and has a higher correlation with FOXP3 and IL2RA expression. While the NIFA Treg factor is significantly negatively associated with response in post-treatment samples, the corresponding human annotation is not (panel A). (C) A new myeloid signature positively associated with response. Heatmap of top loading genes along with the factor values for NIFA factor 24 across all cells identified as “Monocytes/Macrophages” and 1,500 randomly sampled cells. NIFA identifies a subset of the Monocytes/Macrophages cells with unique gene expression pattern. While general myeloid signatures (that is Monocytes/Macrophages and Dendritic cells) were negatively associated with response, the NIFA-24 signature has the opposite pattern (see panel A).

**Table 1.**
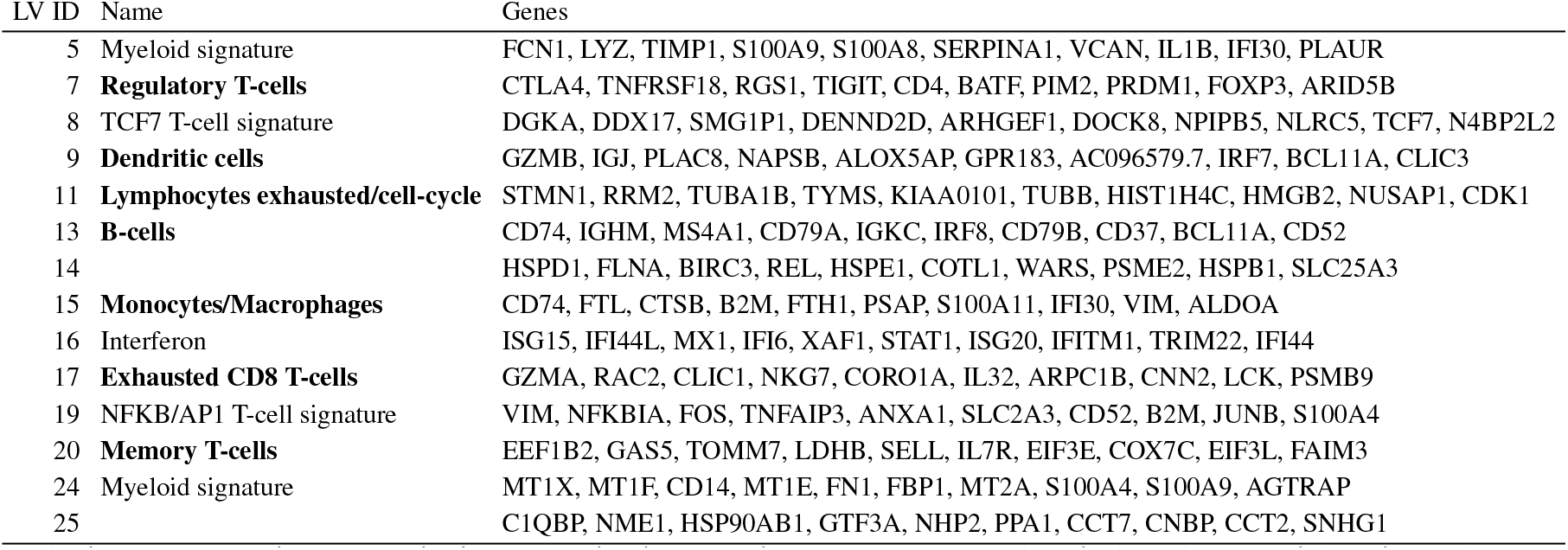
Top loading genes for each factor found to be associated with immunotherapy response. To facilitate biological interpretability, ribosomal genes and genes that we not provided with HGNC symbols were not considered. Factors that best match the known human annotations were named accordingly and are in bold. Other factors could be clearly identified as coherent biological pathways are named based on the loadings. Factor 14 and 25 did not have a clear correspondence to any pathway or cell-type signature and are left unnamed.

Each NIFA factor that has the best correspondence to human annotations is given the same name and the results are grouped with grey ovals in Figure 6. We find that for these matched variables, there is an overall good correspondence between NIFA factors and human-annotated cell types. Specifically, both methods discover B cells as the variable most positively predictive of response and a CD8 T-cell/exhaustion/cell-cycle signature (termed “Lymphocytes exhausted/cell-cycle” in the original study) as the most negatively predictive.

For some subtle patterns the results of NIFA and human annotations can diverge. Human annotations such as “Lymphocytes” and “Cytotoxicity” were not well reproduced by NIFA (correlation of 0.46 and 0.46 respectively) and the corresponding NIFA variables are not significant. On the other hand, NIFA found three different T-cell signatures (8, 19 and 20) which were all associated with the “memory T-cell” human annotation and all were significantly predictive of response. One of these signatures has TCF7 as a top loading gene and thus NIFA was able to independently discover one of the key findings of the original study – that the fraction of TCF7-positive T cells is highly associated with response.

Aside from generally reproducing the main findings of the original study, NIFA was able to uncover additional patterns. For example, we find that presence of Tregs is negatively associated with response in the post-treatment samples. The corresponding human annotation is however not significant despite the fact that the two variables are highly correlated (Pearson correlation = 0.71). Human regulatory T cells are difficult to identify from transcriptional profiles. There are no genes that are *unique* to this cell type. The canonical transcription factor (FOXP3) and surface marker (CD25/IL2RA) can also be transiently expressed by non-Treg CD4 cells (Chen and Oppenheim, 2011); on the other hand, because of noise in scRNA-seq data the absence of these markers doesn’t exclude Treg status. Upon closer inspection, we find that NIFA is more conservative in designating Tregs than the human annotation counterpart. Using the NIFA mixture components we can perform a hard cell-type assignment based on the probability of being in the high-expression component. Using 0.5 as cutoff, NIFA finds 1,418 Tregs, in contrast to 1,740 of human annotated ones. We find that this discrepancy is highly non-random and that the NIFA Tregs are more likely to express both FOXP3 and IL2RA (Figure 6C) indicating that the NIFA Treg signature is more specific.

Overall, within this dataset a large number of the human-annotated cell types and NIFA factors are associated with response but some general patterns emerge. Specifically, the presence of myeloid cell types is negatively associated with response while presence of lymphocytes, exclusive of those with an exhaustion-like phenotype (for example B cells and CD4 memory cells), is positively associated with response (see Figure 6A). The general trend that a high myeloid to lymphocyte ratio is associated with worse outcome is observed across a variety of cancers (Thorsson *et al.,* 2018). Our NIFA-based analysis however finds a myeloid signature (NIFA latent factor 24) that corresponds to a subset of annotated “Monocytes/Macrophages” cells is *positively* associated with response, with an effect size that is similar to the lymphocyte populations (Figure 6A). This myeloid subset is identified by high levels of metallothionein genes (MT1X, MT1F, MT1E and MT2A) and some metabolic genes (see Figure 6B). Metallothioneins are a family of small proteins that play important roles in metal homeostasis and protection against heavy metal toxicity, DNA damage and oxidative stress (Si and Lang, 2018). Their induction in tumor-associated macrophages (TAMs) has been noted (Ge *et al.,* 2012) but to our knowledge this is the first report of an association with clinical outcome.

### Runtime and Scalability

NIFA is a Bayesian method with a large number of parameters and thus is expected to be more computationally demanding than simpler factorization methods such as NMF. However our implementation uses closed-form variational solutions and time-intensive computations are preformed in C++ backend making NIFA competitive with other approaches. Our benchmark dataset based on the 1.3 Million Brain Cell dataset (Zheng *et al.,* 2017) from which we samples 100K cells. Overall we find that NIFA is considerably faster than scCoGAPS and comparable to NMF (see Figure 7 and Supplementary Methods for full details).

**Fig. 7.**
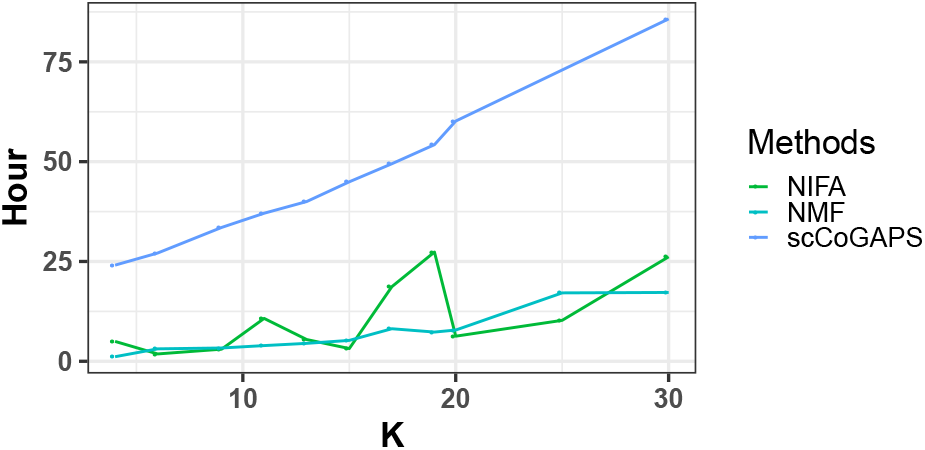
Running time comparison of NIFA, NMF and scCoGAPS with different number of factors *k* on the benchmark dataset.

## 4 Discussion

We propose a factor analysis model designed specifically for single-cell data. The model combines features of PCA, ICA and NMF. Specifically, our model optimizes the PCA-like matrix reconstruction objective with mixture of Gaussian priors on the factors which encourages decomposition along multi-modal directions. We also adopt truncated Gaussian priors on the loadings thus imposing an NMF-like strict non-negativity constraint (see Figure 1E). Using a variational Bayes framework allows us to automatically fit hyperparameters such as the number of Gaussian mixtures. We evaluate our model using both known cell identity and pathway information and demonstrate that NIFA generates biologically coherent factors that align well this prior knowledge.

One additional feature of our model is that this fully Bayesian framework is readily extensible. For example, it easily supports genespecific priors for the loadings. This makes it possible to use known biological pathways as additional constraints. We plan on developing this extension in our future work.

## Acknowledgements

This research was supported in part by the University of Pittsburgh Center for Research Computing through the resources provided.

## Competing interests

The authors declare that they have no competing interests.

## Notes

### Competing Interest Statement

The authors have declared no competing interest.

