## Supplementary Material for "Non-negative Independent Factor Analysis disentangles discrete and continuous sources of variation in scRNA-seq data"

### Supplementary Figures

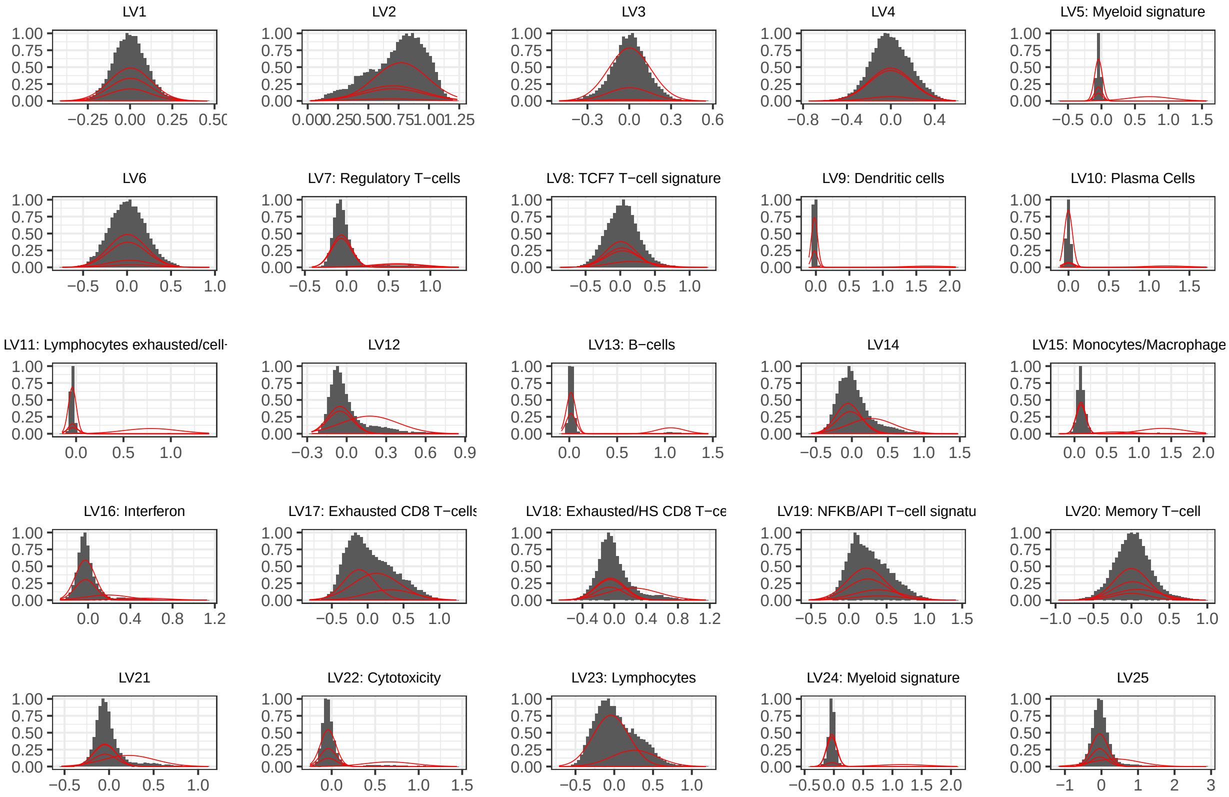

Figure S1: Factor histograms and mixture model fits for the Sade-Feldman *et al.* immunotherapy dataset. Factors that best correspond to cell types identified in the original study are labeled accordingly. NIFA finds both multimodal and unimodal factors and as expected the multimodal factors are more likely to represent cell types

### Supplementary Tables

| Abbreviation | Protocol | Evidence | Type | Tissue | Cells | Cell Types |
|---|---|---|---|---|---|---|
| Camp (Camp *et al.*, 2017) | SMARTer | Homogeneous cell line | Gold | Human: Liver | 777 | 7 |
| ImmunoTherapy (Sade-Feldman *et al.*, 2018) | Smart-Seq2 | k-means clustering | Silver | Human: Metastatic melanoma | 16,291 | 11 |
| Klein (Klein *et al.*, 2015) | inDrop | Principal genes identified by PCA | Silver | Mouse: Embryonic Stem Cells | 2,717 | 4 |
| Kolodziejczyk (Kolodziejczyk *et al.*, 2015) | SMARTer | Homogeneous cell line | Gold | Mouse: Embryonic Stem Cells | 704 | 3 |
| Li (Li *et al.*, 2017) | SMARTer | Homogeneous cell line | Gold | Human: Colorectal Tumors | 561 | 9 |
| Liu (Liu *et al.*, 2019) | 10x drop-seq | k-means clustering & specific markers | Silver | Mouse: Tumor immune cells | 1,607 | 13 |
| Nestorowa (Nestorowa *et al.*, 2016) | Smart-Seq2 | hierachical clustering & specific markers | Silver | Mouse: Hematopoietic Stem Cells | 1,656 | 9 |
| Olsson (Olsson *et al.*, 2016) | SMARTer | Flow cytometry & cell sorting | Gold | Mouse: Hematopoietic Stem Cells | 382 | 4 |
| SimKumar4easy (Duò *et al.*, 2018) | NA | Simulated | Gold | NA | 500 | 4 |
| SimKumar4hard (Duò *et al.*, 2018) | NA | Simulated | Gold | NA | 499 | 4 |
| SimKumar8hard (Duò *et al.*, 2018) | NA | Simulated | Gold | NA | 499 | 8 |
| Zhengmix4eq (Duò *et al.*, 2018) | NA | Simulated | Gold | NA | 3,555 | 4 |
| Zhengmix4uneq (Duò *et al.*, 2018) | NA | Simulated | Gold | NA | 6,414 | 4 |
| Zhengmix8eq (Duò *et al.*, 2018) | NA | Simulated | Gold | NA | 3,971 | 8 |

Table S1: The complete set of datasets evaluated in this study. Gold datasets are those where cell types are determined due to the experimental design (for example by sorting cells). Silver datasets are those where cell types were assigned from the data using biological prior knowledge.

### Supplementary Methods

#### The prior distributions of parameters

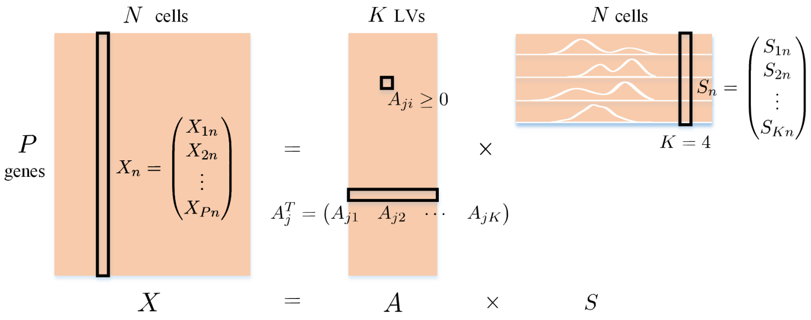

Figure S2: A schematic representation of the NIFA model. $X$ represents a scRNA-seq matrix with dimension $P$-by-$N$, where $P$ is the number of genes and $N$ is the number of cells. Given $X$, we want to infer $A$ which denotes a non-negative loading matrix with dimension $P$-by-$K$ and $S$ which stands for sources or latent variables with dimension $K$-by-$N$. We impose multi-modal priors on the rows of $S$, but the exact number of modes is automatically determined and thus can be one.

The noise parameter $\beta$ comes with a Gamma prior with parameter $a_\beta$ and $b_\beta$.

$$P(\beta) = Gamma(\beta | a_\beta, b_\beta) \tag{1}$$

The membership indicator $\epsilon_{imn}$ comes with a Bernoulli prior with mixing proportion $\pi_{im}$.

$$P(\epsilon_{imn}) = \pi_{im}^{\epsilon_{imn}} \tag{2}$$

The $\mu_{im}$ is assumed to follow a Gaussian distribution with $\rho_{im}$ as the mean and $\phi_{im}$ as the inverse of the variance.

$$P(\mu_{im}) = N(\mu_{im} | \rho_{im}, \phi_{im}) \tag{3}$$

The inverse of the variance $\sigma$ follows a Gamma distribution with parameters $a_{\sigma_{im}}$ and $b_{\sigma_{im}}$.

$$P(\sigma_{im}) = Gamma(\sigma_{im} | a_{\sigma_{im}}, b_{\sigma_{im}}) \tag{4}$$

The dependency structure of the NIFA model is summarized in Figure S3.

#### Variational update functions

Here we provide detailed derivation for $q(A_{ji})$ and list update functions for other $q(\cdot)$.

##### Derive the update equation for $q(A_{ji})$

$$q(A_{ji}) = f(A_{ji} | \eta_{ji}^*, \lambda_i^*, 0, \infty) \tag{5}$$

which is the truncated normal distribution with $a = 0$ and $b = \infty$. where

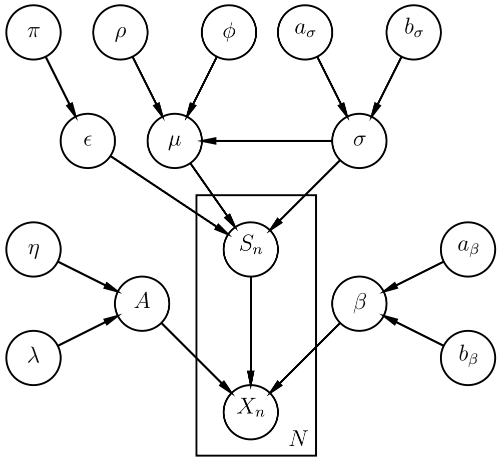

Figure S3: Parameters of the NIFA model are summarized in a directed acyclic graph.

$$\begin{cases} \lambda_i^* = \lambda_i + < \beta > \sum_{n=1}^{N} < S_{in}^2 > \\ \eta_{ji}^* = \frac{\lambda_i \eta_{ji} - <\beta> \left[ \sum_{n=1}^{N} <S_{in}> (\sum_{m \neq i} <A_{jm}><S_{mn}> - X_{jn}) \right]}{\lambda_i + <\beta> \sum_{n=1}^{N} <S_{in}^2>} \end{cases}$$

Derivation starts here. We first derive the formula of $\log P(A_{ji}|\eta_{ji}, \lambda_i)$ which will be used in the next step.

$$\begin{aligned} & \log P(A_{ji}|\eta_{ji}, \lambda_i) \\ = & -\frac{1}{2}\log(2\pi) + \frac{1}{2}\log\lambda_i - \frac{1}{2}\lambda_i(A_{ji} - \eta_{ji})^2 - \log(1 - \Phi(-\eta_{ji}\lambda_i^{\frac{1}{2}})) \\ \xrightarrow{\text{remove constant}} & -\frac{1}{2}\lambda_i(A_{ji}^2 - 2A_{ji}\eta_{ji}) \end{aligned}$$

We are going to find the $q$ density for $A_{ji}$ and we start from $\prod_{n=1}^{N} P(X_n|A, S_n, \beta)P(A_{ji}|\eta_{ji}, \lambda_i)$.

$$\prod_{n=1}^{N}(2\pi)^{-\frac{P}{2}}\beta^{\frac{P}{2}}\exp\left(-\frac{\beta}{2}\sum_{j=1}^{P}(X_{jn}-A_j^T S_n)^2\right)\cdot P(A_{ji}|\eta_{ji},\lambda_i)$$
$$=\prod_{n=1}^{N}(2\pi)^{-\frac{P}{2}}\beta^{\frac{P}{2}}\exp\left(-\frac{\beta}{2}\sum_{j=1}^{P}(A_{ji}S_{in}+\sum_{m\neq i}A_{jm}S_{mn}-X_{jn})^2\right)\cdot P(A_{ji}|\eta_{ji},\lambda_i)$$
$$\xRightarrow{\log}\sum_{n=1}^{N}\left[-\frac{P}{2}\log(2\pi)+\frac{P}{2}\log\beta-\frac{\beta}{2}\sum_{j=1}^{P}(A_{ji}S_{in}+\sum_{m\neq i}A_{jm}S_{mn}-X_{jn})^2\right]+\log P(A_{ji}|\eta_{ji},\lambda_i)$$
$$\xRightarrow{\text{remove constant}}\sum_{n=1}^{N}\left[-\frac{\beta}{2}(A_{ji}S_{in}+\sum_{m\neq i}A_{jm}S_{mn}-X_{jn})^2\right]+\log P(A_{ji}|\eta_{ji},\lambda_i)$$
$$\xRightarrow{\text{remove constant}}\sum_{n=1}^{N}\left[-\frac{\beta}{2}\left(A_{ji}^2S_{in}^2+2A_{ji}S_{in}(\sum_{m\neq i}A_{jm}S_{mn}-X_{jn})\right)\right]+\log P(A_{ji}|\eta_{ji},\lambda_i)$$
$$=-\frac{\beta}{2}\left[A_{ji}^2\sum_{n=1}^{N}S_{in}^2+2A_{ji}\sum_{n=1}^{N}S_{in}(\sum_{m\neq i}A_{jm}S_{mn}-X_{jn})\right]-\frac{1}{2}\lambda_i(A_{ji}^2-2A_{ji}\eta_{ji})$$
$$=A_{ji}^2\left(-\frac{\beta}{2}\sum_{n=1}^{N}S_{in}^2-\frac{1}{2}\lambda_i\right)+A_{ji}\left[-\beta\sum_{n=1}^{N}S_{in}(\sum_{m\neq i}A_{jm}S_{mn}-X_{jn})+\eta_{ji}\lambda_i\right]$$
$$\xRightarrow{<\cdot>_Q}A_{ji}^2(-\frac{<\beta>}{2}\sum_{n=1}^{N}<S_{in}>^2-\frac{1}{2}\lambda_i)+A_{ji}\left[-<\beta>\sum_{n=1}^{N}<S_{in}>(\sum_{m\neq i}<A_{jm}><S_{mn}>-X_{jn})+\eta_{ji}\lambda_i\right]$$

We can recognize this as a Gaussian distribution. The mean and variance parameters for $q(A_{ji})$ are as follows,

$$\begin{cases}\lambda_i^*=\lambda_i+<\beta>\sum_{n=1}^{N}<S_{in}^2>\\ \eta_{ji}^*=\frac{\lambda_i\eta_{ji}-<\beta>\left[\sum_{n=1}^{N}<S_{in}>(\sum_{m\neq i}<A_{jm}><S_{mn}>-X_{jn})\right]}{\lambda_i+<\beta>\sum_{n=1}^{N}<S_{in}^2>}\end{cases}$$

**Other update functions** $q(\cdot)$

**Update equation** $q(S_n)$

$$q(S_n)=N(S_n|\mu_m^*,\Sigma_m^*) \tag{6}$$

where

$$\begin{cases}\mu_m^*=\Sigma_m^*\left[\sum_{m=1}^{M}B_m^{-1}\mu_m+<\beta>\sum_{j=1}^{P}X_{jn}<A_j>\right]\\ \Sigma_m^*=\left[\sum_{m=1}^{M}B_m^{-1}+<\beta>\sum_{j=1}^{P}<A_jA_j^T>\right]^{-1}\end{cases}$$

$$\mu_m=\begin{pmatrix}<\mu_{1m}>\\<\mu_{2m}>\\ \vdots\\<\mu_{Km}>\end{pmatrix}, B_m^{-1}=\begin{pmatrix}<\epsilon_{1mn}><\sigma_{1m}> & & \\ & \ddots & \\ & & <\epsilon_{Kmn}><\sigma_{Km}>\end{pmatrix}$$

**Update equation** $q(\epsilon_{imn})$

$$q(\epsilon_{imn}) = \widetilde{\lambda_{imn}}^{\epsilon_{imn}} \tag{7}$$

where

$$\widetilde{\lambda_{imn}} = \frac{e^{\lambda_{imn}}}{\sum_{m=1}^{M} e^{\lambda_{imn}}}$$
$$\lambda_{imn} = -\frac{1}{2}\log(2\pi) + \frac{1}{2} < \log\sigma_{im} > + \log\pi_{im} - \frac{1}{2} < \sigma_{im} > \left[< S_{in}^2 > -2 < S_{in} >< \mu_{im} > + < \mu_{im}^2 >\right]$$

**Update equation** $q(\mu_{im})$

$$q(\mu_{im}) = N(\mu_{im}|\rho_{im}^*, \phi_{im}^*) \tag{8}$$

where

$$\begin{cases} \rho_{im}^* = (\phi_{im}^* < \sigma_{im} >)^{-1} \left[< \sigma_{im} > \sum_{n=1}^{N} < \epsilon_{imn} >< S_{in} > + \phi_{im}\rho_{im} < \sigma_{im} >\right] \\ \phi_{im}^* = \phi_{im} + \sum_{n=1}^{N} < \epsilon_{imn} > \end{cases}$$

**Update equation** $q(\sigma_{im})$

$$q(\sigma_{im}) = Gamma(\sigma_{im}|a_{\sigma_{im}}^*, b_{\sigma_{im}}^*) \tag{9}$$

where

$$\begin{cases} a_{\sigma_{im}}^* = a_{\sigma_{im}} + \frac{1}{2}\sum_{n=1}^{N} < \epsilon_{imn} > + \frac{1}{2} \\ b_{\sigma_{im}}^* = b_{\sigma_{im}} + \dfrac{1}{2}\displaystyle\sum_{n=1}^{N} < \epsilon_{imn} > (< S_{in}^2 > -2 < S_{in} >< \mu_{im} > + < \mu_{im}^2 >) \\ \qquad + \dfrac{1}{2} < \phi_{im} > (< \mu_{im}^2 > -2 < \mu_{im} > \rho_{im} + \rho_{im}^2) \end{cases}$$

**Update equation** $q(\beta)$

$$q(\beta) = Gamma(\beta|a_\beta^*, b_\beta^*) \tag{10}$$

where

$$\begin{cases} a_\beta^* = a_\beta + \frac{NP}{2} \\ b_\beta^* = b_\beta + \frac{1}{2}\sum_{n=1}^{N}\sum_{j=1}^{P}[X_{in}^2 - 2X_{in} < A_j^T >< S_n > + Tr(< A_j A_j^T >< S_n S_n^T >)] \end{cases}$$

**Update equation** $q(\pi_{im})$

$$\pi_{im} = \frac{1}{N}\sum_{n=1}^{N}\widetilde{\lambda_{imn}} \tag{11}$$

**Update functions of expectations**

$$< S_n >= \mu_m^* \tag{12}$$

$$< S_n S_n^T >= \mu_m^* {\mu_m^*}^T + \Sigma_m^* \tag{13}$$

$$< \epsilon_{imn} >= \widetilde{\lambda_{imn}} \tag{14}$$

$$< \sigma_{im} >= \frac{a_{\sigma_{im}}^*}{b_{\sigma_{im}}^*} \tag{15}$$

$$< A_j >= \mu_j \tag{16}$$

$$< A_j A_j^T >= \mu_j \mu_j^T + \Sigma_j \tag{17}$$

$$< \log \sigma_{im} >= \psi(a^*_{\sigma_{im}}) - \ln(b^*_{\sigma_{im}}) \tag{18}$$

$$< \mu_{im} >= \rho^*_{im} \tag{19}$$

$$< \mu^2_{im} >= (\phi^*_{im}\sigma^*_{im})^{-1} + (\rho^*_{im})^2 \tag{20}$$

$$< \beta >= \frac{a^*_\beta}{b^*_\beta} \tag{21}$$

**Update functions of $\mu_j$ and $\Sigma_j$ for Eq. 16 and Eq. 17**

$$\delta_i = \sqrt{(\lambda^*_i)^{-1}}$$

$$Z_{ji} = 1 - \Phi(-\frac{\eta^*_{ji}}{\delta_i})$$

$$\mu_j \quad \Leftarrow \quad \mu_{ji} = \eta^*_{ji} + \frac{\Phi(-\frac{\eta^*_{ji}}{\delta_i})}{Z_{ji}}\delta_i \tag{22}$$

$$\Sigma_j \quad \Leftarrow \quad \Sigma^{ii}_j = \delta^2 \left[ 1 + \frac{-\frac{\eta^*_{ji}}{\delta_i}\Phi(-\frac{\eta^*_{ji}}{\delta_i})}{Z} - \left( \frac{\Phi(-\frac{\eta^*_{ji}}{\delta_i})}{Z} \right)^2 \right] \tag{23}$$

where $\Sigma_j$ is a diagonal matrix and $\Sigma^{ii}_j$ stands for the $i$th diagonal values of $\Sigma_j$.

#### Preprocessing pipeline

The data preprocessing pipeline is illustrated in Figure S4. Some pre-processing steps were only applied to certain methods. For example, we employed SVD smoothing (that is reconstructing the input as a truncated SVD with rank=50) because it makes the constant-variance Gaussian error assumption more valid for NIFA. However, we found that empirically this had little effect on the results. For NMF we used KL divergence (or equivalently a Poisson error model) which is most appropriate for unsmoothed data. For NIFA we chose to z-score the input by row (gene). The z-scoring operation theoretically makes it easier to pick up on small-variance but highly deferentially expressed genes and it produced modest improvement in most (though not all) benchmark datasets. Row z-scoring was not applied to NMF as it produces negative numbers. It was also not applied to ICA as the ICA objective function references only the shape of the factor distribution and thus is invariant under row scaling.

#### NIFA initialization

NIFA updates are relatively expensive and thus a good initialization can significantly improve running time. We initialize NIFA by a simple matrix decomposition with non-negativity constraint on the loadings, which is formulated by Eq. 24. This problem can be efficiently solved by the alternating least squares algorithm.

$$\begin{gathered} \min_{A,S} ||X - AS||_F + \lambda_1 ||A||_F + \lambda_2 ||S||_F \\ \text{subject to } A \geq 0 \end{gathered} \tag{24}$$

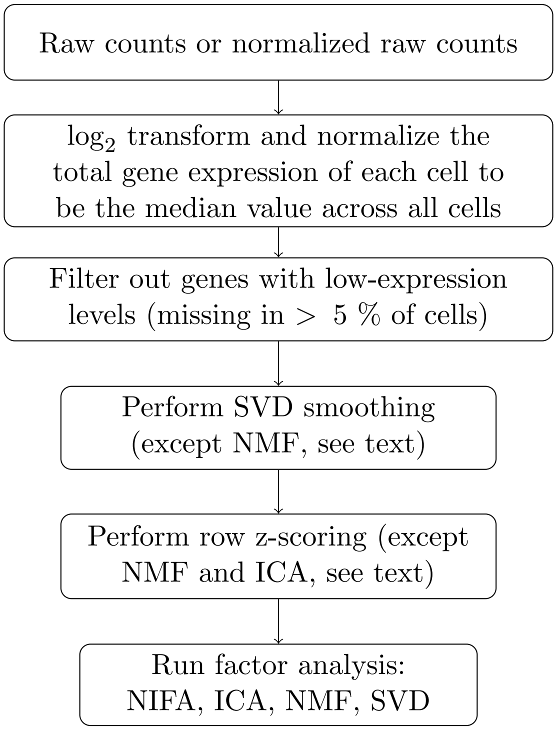

Figure S4: Preprocessing workflow.

The key to good initialization is selecting $\lambda_1$ and $\lambda_2$ so that the residual ($E((X - AS)^2)$) is close to what one would get with a Bayesian framework. We find that a simple heuristic works well in this regard. Given the truncated rank-$k$ SVD of the data $UDV^T$ we set $\lambda_2 = D_k$ and $\lambda_1 = \lambda_2/2$.

#### Simulation details

The dimension of the simulated matrix $X$ is set to be 2000-by-500 (gene-by-sample). There are 6 latent factors, each of which contains 2 Gaussian mixtures. The dimension of loading matrix $A$ is 2000-by-6 with each column corresponding to a single loading and the matrix $S$ is 6-by-500 with each row corresponding to a single latent factor. We draw the first loading vector from Gamma distribution $\Gamma(5, 1)$. Then the subsequent loadings are simulated by adding noise following Gaussian distribution $N(0, 2)$ to the first simulated loading. We take the absolute values of noise to make sure loadings are kept positive. In this way, we can control the collinearity to simulate correlated loadings. Each latent factor is generated in a hierarchical manner. First we draw mean and variance parameters for the first Gaussian mixture associated with each latent factor from Gaussian distribution $N(2, 10)$ and Gamma distribution $\Gamma(10, 1)$ correspondingly. Regarding each latent factor the rest of mixtures are generated by adding noise drawn from a uniform distribution $U(2, 4)$ to the mean of the first mixture. Then each entry in the latent factor is assigned to any of the mixtures with probability and the exact value is drawn based on the distribution of assigned mixture. Finally $X$ is generated as the sum of $AS$ and noise drawn from Gaussian distribution $N(0, 0.1)$. In order to generate non-negative input to NMF, we offset the matrix by a minimum constant $c = \min(C)$ which makes the result $X + c$ non-negative.

##### Evaluation details

We compare NIFA with NMF (Gaujoux, 2018), ICA (fastICA implementation (Marchini *et al.*, 2019)) and scCoGAPS (Stein-O'Brien *et al.*, 2019). For NMF we used KL loss which we found dramatically outperformed the square loss alternative.

***Cell-Type Correspondence*** We compute the absolute Pearson correlations between each pair of cell type (binary indicator) and decomposed factor. Each cell type annotation is assigned to the factor that is best correlated with the corresponding cell-type binary indicator.

***Pathway Enrichment*** We perform a hypergeometric test on each of the loadings with the top 500 genes as foreground and the rest as background. For SVD and ICA, we use the absolute values of loadings to perform this enrichment test. The $p$ values are adjusted with Benjamini-Hochberg procedure and we denote pathways with adjusted $p$-val $< 0.05$ as significantly enriched pathways. The pathway enrichment is summarized as average fold enrichment across the significant loading-pathway associations.

##### Running Time Analysis

We compare the running time of NIFA to that of NMF (nnls R package, C++ backend) (Lin and Boutros, 2019) and scCoGAPS which is a Bayesian NMF approach that is solved via MCMC (Fertig *et al.*, 2010). We used KL-divergence as the objective for timing NMF. This objective was also used for our interpretability comparisons as it produces a dramatic improvement in performance over the faster least squares objective. Since the methods compared differ in parallelization, hyper-parameter settings, and convergence criteria, the specific settings used to conduct the experiment are summarized in Table.S2.

| Methods | Versions | Convergence criteria | Maximum iterations | Number of cores/threads |
|---|---|---|---|---|
| NIFA | 0.1.0 | Stop when the relative changes on the latent variable matrix less or equal to 5e-5 | 1000 (number of steps of updating all variational parameters) | 1 thread |
| scCoGAPS | 3.4.1 | NA | 1000 (number of iterations for each phase of algorithm) | 2 cores, 1 thread/core |
| NNLM | 0.4.3 | relative tolerance is 1e-4 | 500 (number of iterations of alternating NNLS solutions to H and W) | 8 threads |

Table S2: Method specific settings for timing experiments.
